# Neuronal cascades shape whole-brain functional dynamics at rest

**DOI:** 10.1101/2020.12.25.424385

**Authors:** Giovanni Rabuffo, Jan Fousek, Christophe Bernard, Viktor Jirsa

## Abstract

At rest, mammalian brains display remarkable spatiotemporal complexity, evolving through recurrent brain states on a slow timescale of the order of tens of seconds. While the phenomenology of the resting state dynamics is valuable in distinguishing healthy and pathological brains, little is known about its underlying mechanisms. Here, we identify neuronal cascades as a potential mechanism. Using full-brain network modeling, we show that neuronal populations, coupled via a detailed structural connectome, give rise to large-scale cascades of firing rate fluctuations evolving at the same time scale of resting-state networks. The ignition and subsequent propagation of cascades depend upon the brain state and connectivity of each region. The largest cascades produce bursts of Blood-Oxygen-Level-Dependent (BOLD) co-fluctuations at pairs of regions across the brain, which shape the simulated resting-state network dynamics.We experimentally confirm these theoretical predictions. We demonstrate the existence and stability of intermittent epochs of functional connectivity comprising BOLD co-activation bursts in mice and human fMRI. We then provide evidence for the existence and leading role of the neuronal cascades in humans with simultaneous EEG/fMRI recordings. These results show that neuronal cascades are a major determinant of spontaneous fluctuations in brain dynamics at rest.

**Significance Statement:** Functional connectivity and its dynamics are widely used as a proxy of brain function and dysfunction. Their neuronal underpinnings remain unclear. Using connectome-based modeling, we link the fast microscopic neuronal scale to the slow emergent whole-brain dynamics. We show that cascades of neuronal activations spontaneously propagate in resting state-like conditions. The largest neuronal cascades result in the co-fluctuation of Blood-Oxygen-Level-Dependent signals at pairs of brain regions, which in turn translate to stable brain states. Thus, we provide a theoretical framework for the emergence and the dynamics of resting-state networks. We verify these predictions in empirical mouse fMRI and human EEG/fMRI datasets measured in resting states conditions. Our work sheds light on the multiscale mechanisms of brain function.

## 2 Introduction

At rest, functional Magnetic Resonance Imaging (fMRI) reveals the existence of periods during which blood-oxygenation-level-dependent (BOLD) activity is highly correlated between specific brain regions, known as resting-state networks (*RSNs*). *RSNs* are consistently observed across several mammalian species including humans (Fox & Raichle, 2007; Power et al., 2011) and non-human (Vincent et al., 2007) primates, as well as in rats (Lu et al., 2012; Upadhyay et al., 2011) and mice (Gozzi & Schwarz, 2016; Grandjean et al., 2020; Sforazzini, Schwarz, Galbusera, Bifone, & Gozzi, 2014; Stafford et al., 2014). Functional Connectivity (*FC*) can be used to characterize these strongly correlated functional communities of brain network nodes. A growing body of research on mammalian species emphasizes the dynamic nature of *RSNs*, showing that large-scale *FC* patterns switch between stable and unstable epochs at an infraslow pace (*<* 0.1Hz) (Allen et al., 2014; Gonzalez-Castillo & Bandettini, 2018; Grandjean et al., 2017; Hindriks et al., 2016; Preti & Van De Ville, 2017; Qin et al., 2015; Tagliazucchi, Von Wegner, Morzelewski, Brodbeck, & Laufs, 2012). Dynamic *FC* (*dFC*; (Hutchison et al., 2013)) keeps track of *FC* fluctuations and provides a marker of healthy, ageing, and diseased brains (Battaglia et al., 2020; Damaraju et al., 2014; Du et al., 2017; Preti, Bolton, & Van De Ville, 2017; Rashid, Damaraju, Pearlson, & Calhoun, 2014; Su et al., 2016). Our current understanding of these phenomena rests upon computational studies, performed at the whole-brain level, suggesting that the *RSN* dynamics is an emergent property of the network (Ponce-Alvarez et al., 2015), which operates near criticality (Deco & Jirsa, 2012; Ghosh, Rho, McIntosh, Kötter, & Jirsa, 2008; Jirsa et al., 2017). Nevertheless, the range of possible mechanistic underpinnings remains vast and requires further narrowing down. Here we address this issue and investigate *in silico* the potential neuronal mechanisms giving rise to whole-brain network dynamics, generate predictions and test them *in vivo*. We use The Virtual Brain (TVB), a neuroinformatics and simulation platform, which allows connectome-based whole-brain modeling of multiple species including humans (Sanz-Leon, Knock, Spiegler, & Jirsa, 2015) and mice (Melozzi, Woodman, Jirsa, & Bernard, 2017). TVB includes a fixed number of network nodes (at least one per brain region) and the physical connections that link them, such as white matter tracts defined by diffusion tensor imaging or axon projections obtained with virus injections (Sanz Leon et al., 2013). A large number of neural mass models are readily available in TVB, which produce activity of neuronal populations and map on a range of brain imaging modalities including BOLD fMRI, EEG, and MEG signals. We here adapt a novel neural mass model (NMM), which has been previously derived as the exact limit of an infinite number of all-to-all coupled quadratic integrate-and-fire neurons (Montbrío, Paźo, & Roxin, 2015) (Figure 1a). This analytic step allows to derive the average firing rate and membrane potential of a mesoscopic neuronal population, thus providing suitable neural mass variables while keeping track of the internal spiking neural network organization, which is important to infer potential mechanisms at the neuronal scale. Using this NMM, we report a mechanism by which spontaneous local re-organizations of regional spiking neural net-works, can trigger global infra-slow fluctuations, which we name neuronal cascades. The largest neuronal cascades give rise to bursts of simulated BOLD co-activations (BOLD-CA) at pairs of brain regions across the brain, which in turn account for stable long-lasting *FC* states and their dynamics. We verify experimentally the link between BOLD-CA and *dFC* in mouse fMRI and human EEG-fMRI. On the simultaneous human EEG acquisition, we discover the presence of neuronal cascades and we demonstrate their role in driving the switching behavior of *FC* states. These findings provide the first evidence of a multiscale mechanism underlying resting-state network dynamics which can be robustly assessed in non-invasive brain imaging signals.

**Figure 1:**
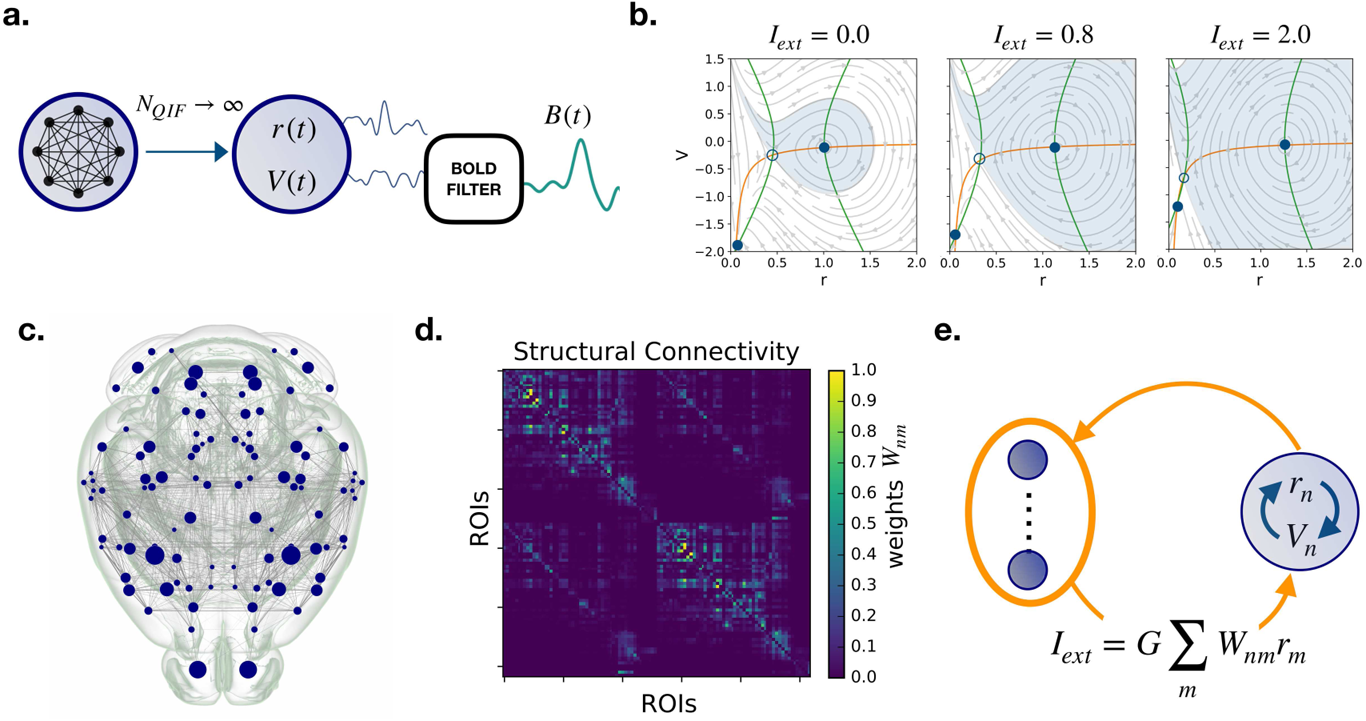
Connectome based modeling. (a) The mean firing rate *r* and membrane potential *V* variables of the Neural Mass Model (NMM) are derived as the limit of infinite all-to-all coupled quadratic integrate-and-fire (QIF) neurons. Applying the Balloon-Windkessel model to *V_n_*(*t*) we obtain the simulated BOLD signal *B_n_*(*t*). (b) The phase plane of each decoupled node (*I_ext_* = 0) has a ‘down’ stable fixed point and an ‘up’ stable focus (full dots). These points are defined at the intersection of the nullclines 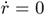 (orange line) and 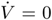 (green line) where the dynamics freezes. The empty circle marks an unstable fixed point. As the external current *I_ext_* is increased, the phase plane of the neural mass changes (see equations in Methods). In particular, the basin of attraction of the up state gradually becomes larger than that of the down state, while the fixed points move farther apart. (c-d) The mouse connectome and structural connectivity *W_nm_* were imported from the tracer experiments of the Allen Institute. The 104 cortical regions of interest (corresponding to network nodes) are specified in Table 1. (e) When the regions are coupled in a brain network, each node *n* receives an input current *I_ext_* which is the sum of the other nodes’ firing rates, weighted by the structural connectivity. According to panel (b), this input provokes a distortion of the local phase plane at node *n*.

**Table 1:** List of brain regions of interest of the Allen Mouse Atlas considered in the simulations.

|  |  |  |  |
| --- | --- | --- | --- |
| 0 | Right Primary motor area | 52 | Left Primary motor area |
| 1 | Right Secondary motor area | 53 | Left Secondary motor area |
| 2 | Right Primary somatosensory area, nose | 54 | Left Primary somatosensory area, nose |
| 3 | Right Primary somatosensory area, barrel field | 55 | Left Primary somatosensory area, barrel field |
| 4 | Right Primary somatosensory area, lower limb | 56 | Left Primary somatosensory area, lower limb |
| 5 | Right Primary somatosensory area, mouth | 57 | Left Primary somatosensory area, mouth |
| 6 | Right Primary somatosensory area, upper limb | 58 | Left Primary somatosensory area, upper limb |
| 7 | Right Supplemental somatosensory area | 59 | Left Supplemental somatosensory area |
| 8 | Right Gustatory areas | 60 | Left Gustatory areas |
| 9 | Right Visceral area | 61 | Left Visceral area |
| 10 | Right Dorsal auditory area | 62 | Left Dorsal auditory area |
| 11 | Right Primary auditory area | 63 | Left Primary auditory area |
| 12 | Right Ventral auditory area | 64 | Left Ventral auditory area |
| 13 | Right Primary visual area | 65 | Left Primary visual area |
| 14 | Right Anterior cingulate area, dorsal part | 66 | Left Anterior cingulate area, dorsal part |
| 15 | Right Anterior cingulate area, ventral part | 67 | Left Anterior cingulate area, ventral part |
| 16 | Right Agranular insular area, dorsal part | 68 | Left Agranular insular area, dorsal part |
| 17 | Right Retrosplenial area, dorsal part | 69 | Left Retrosplenial area, dorsal part |
| 18 | Right Retrosplenial area, ventral part | 70 | Left Retrosplenial area, ventral part |
| 19 | Right Temporal association areas | 71 | Left Temporal association areas |
| 20 | Right Perirhinal area | 72 | Left Perirhinal area |
| 21 | Right Ectorhinal area | 73 | Left Ectorhinal area |
| 22 | Right Main olfactory bulb | 74 | Left Main olfactory bulb |
| 23 | Right Anterior olfactory nucleus | 75 | Left Anterior olfactory nucleus |
| 24 | Right Piriform area | 76 | Left Piriform area |
| 25 | Right Cortical amygdalar area, posterior part | 77 | Left Cortical amygdalar area, posterior part |
| 26 | Right Field CA1 | 78 | Left Field CA1 |
| 27 | Right Field CA3 | 79 | Left Field CA3 |
| 28 | Right Dentate gyrus | 80 | Left Dentate gyrus |
| 29 | Right Entorhinal area, lateral part | 81 | Left Entorhinal area, lateral part |
| 30 | Right Entorhinal area, medial part, dorsal zone | 82 | Left Entorhinal area, medial part, dorsal zone |
| 31 | Right Subiculum | 83 | Left Subiculum |
| 32 | Right Caudoputamen | 84 | Left Caudoputamen |
| 33 | Right Nucleus accumbens | 85 | Left Nucleus accumbens |
| 34 | Right Olfactory tubercle | 86 | Left Olfactory tubercle |
| 35 | Right Substantia innominata | 87 | Left Substantia innominata |
| 36 | Right Lateral hypothalamic area | 88 | Left Lateral hypothalamic area |
| 37 | Right Superior colliculus, sensory related | 89 | Left Superior colliculus, sensory related |
| 38 | Right Inferior colliculus | 90 | Left Inferior colliculus |
| 39 | Right Midbrain reticular nucleus | 91 | Left Midbrain reticular nucleus |
| 40 | Right Superior colliculus, motor related | 92 | Left Superior colliculus, motor related |
| 41 | Right Periaqueductal gray | 93 | Left Periaqueductal gray |
| 42 | Right Pontine reticular nucleus, caudal part | 94 | Left Pontine reticular nucleus, caudal part |
| 43 | Right Pontine reticular nucleus | 95 | Left Pontine reticular nucleus |
| 44 | Right Intermediate reticular nucleus | 96 | Left Intermediate reticular nucleus |
| 45 | Right Central lobule | 97 | Left Central lobule |
| 46 | Right Culmen | 98 | Left Culmen |
| 47 | Right Simple lobule | 99 | Left Simple lobule |
| 48 | Right Ansiform lobule | 100 | Left Ansiform lobule |
| 49 | Right Paramedian lobule | 101 | Left Paramedian lobule |
| 50 | Right Copula pyramidis | 102 | Left Copula pyramidis |
| 51 | Right Paraflocculus | 103 | Left Paraflocculus |

## 3 Results

We start by introducing the synthetic setup used to simulate the main feature of resting-state network dynamics i.e., the intermittent epochs of stable *FC*. To achieve recurrent functional network activation we set the brain regions into a bistable regime, where the mean membrane potentials can exhibit a resting (’down’) and a depolarized (’up’) state (Hansen, Battaglia, Spiegler, Deco, & Jirsa, 2015). When the regions are suitably coupled over a connectome, the local activity spontaneously fluctuates between these states, which promotes system metastability i.e., the recurrent exploration of multiple network configurations (Deco, Kringelbach, Jirsa, & Ritter, 2017; Jimenez-Marin et al., 2019). In order to understand the mechanisms allowing the dynamic coordination of bistable neural masses into multistable network re-configurations, we looked in detail at the building blocks of our simulation.

### 3.1 Rules of single and coupled nodes dynamics

Let us first consider the dynamics of a single isolated brain node (not connected to the network) where the local NMM parameters are tuned to ensure bistability. If there is no input current (*I_ext_* = 0), solving the model equations (see Methods) resolves the dynamics of the node in terms of the mean firing rate *r*(*t*) and mean membrane potential *V* (*t*). The motion trajectories are represented on the (*r, V*) phase plane (left panel in Figure 1b).

If, at a given time, a node takes a given (*r^★^*, *V^★^*) value, the direction of the arrow at these coordinates indicates the direction of the flow, and where the node dynamics will end up: either falling in the down stable state where the neurons populating the NMM exhibit low firing rate and strong (negative) membrane potential; or spiraling in a damped oscillation into the up stable state, in which the neurons display high average firing rate and low average membrane potential. Thus, in the absence of noise, neuronal activity of the node would rapidly freeze into a stable up or down level of activity (full dots in Figure 1b).

Introducing noise allows the stochastic exploration of the phase space. If the noise is sufficiently large, a random movement in the phase plane can provoke the sudden switch from up→down or down→up basins of attraction (Figure 1b, light blue and white shades respectively), therefore changing the *r* and *V* dynamic modality.

We now consider a non-null input current *I_ext_*, simulating a constant synaptic input. As *I_ext_* increases (Figure 1b), the two branches of the *V*-nullcline 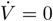 lines) move horizontally away from each other, while the *r*-nullcline (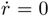 line) remains fixed. Correspondingly, the location of the stable points in the phase plane shifts (at 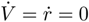) and their basins of attraction change in size. The separatrix i.e., the line separating the up and down basins of attraction, is therefore a dynamical element: its location depends on the amount of input that the node is receiving at a given time.

If *I_ext_* is small, the separatrix is close to the up state (Figure 1b, left) and the crossing of the separatrix via random fluctuations is more probable in the up → down direction; the node explores only the down state. Conversely, if the separatrix is close to the down state when *I_ext_* is high (Figure 1b, right), the node likely remains in the up state.

This property of the NMM is key to understand the systems dynamic. At a given time *t* the inputs received by node *n* will shift the separatrix, and make it easier (or not) for the node to change its activity as a function of noise and of its current state ((*r^★^*, *V ^★^*) coordinates at time *t*).

Using this NMM, we simulate whole-brain dynamics in a mouse brain avatar (Melozzi et al., 2019, 2017), which detailed connectome is imported from the tracer experiments of the Allen Institute and is parcellated into anatomic regions of interest, associated with the network nodes (Figure 1c) (Oh et al., 2014). The communication between regions is weighted by the structural links *W_nm_* of the structural connectivity (Figure 1d). Thus, the input *I_ext_* of a node represents the synaptic drive from the firing rates *r_m_* of all the *m* nodes that are connected to node *n* (Figure 1e; see Methods Section). As an effect of the coupling, the (projected) phase plane (*r_n_, V_n_*) at node *n* can be thought of as a distorted version of the phase plane of a single isolated node, where the location of the separatrix depends on the structural role that region *n* plays in the network hierarchy. Less connected regions receive a weak input current *I_n_* so that the local phase plane is slightly distorted. Instead, regions with high centrality in the connectome, or that are part of strongly connected structural motifs, receive a strong input, which causes a greater distortion of the local phase plane, with consequences on their activity.

### 3.2 Co-activation bursts account for RSN dynamics

Once a working regime is chosen by the selection of global and local model parameters, the raw outcome of the simulation consists of a high time-resolution neuroelectric signal: the average firing rate *r* and membrane potential *V* for each node. As a convention, we refer to the average firing rate activity as the simulated EEG data. With a further processing step, we obtain a low time-resolution simulated BOLD activity by filtering the membrane potentials through the Balloon-Windkessel model (Friston, Mechelli, Turner, & Price, 2000) (Figure 1a, right).

In order to compare simulated and empirical whole-brain imaging data, we analyze the dynamic evolution of functional brain patterns with *dFC* measures (Hutchison et al., 2013; Karahanoğlu & Van De Ville, 2015; Lindquist, Xu, Nebel, & Caffo, 2014; Majeed et al., 2011; Shine et al., 2015; Smith et al., 2012). The most diffuse definition of *dFC*, is based on a sliding window approach (Allen et al., 2014). In brief, inside each time window, a static *FC* is computed as the correlation matrix of the BOLD activities. Then, the entries of the windowed-*dFC* (*dFC_w_*) matrix are defined as the correlation between the *FC*s at different windows (see Methods). In typical empiric datasets, *dFC_w_* matrices show non-trivial block structures; diagonal block structures represent epochs of stable *FC*, while the off-diagonal blocks mark the re-occurrence of correlated *FC* at distinct times. For our analysis, we introduce a *dFC* measure derived in an edge-centric approach, where the dynamics for the edge *E_nm_* is defined by the product of the z-scored BOLD activities *B_n_* and *B_m_* at nodes *n* and *m* (Faskowitz, Esfahlani, Jo, Sporns, & Betzel, 2020). The resulting edge co-activation (CA) time series (Figure 2a, orange box) tracks the temporal unfold of correlations across node pairs.

**Figure 2:**
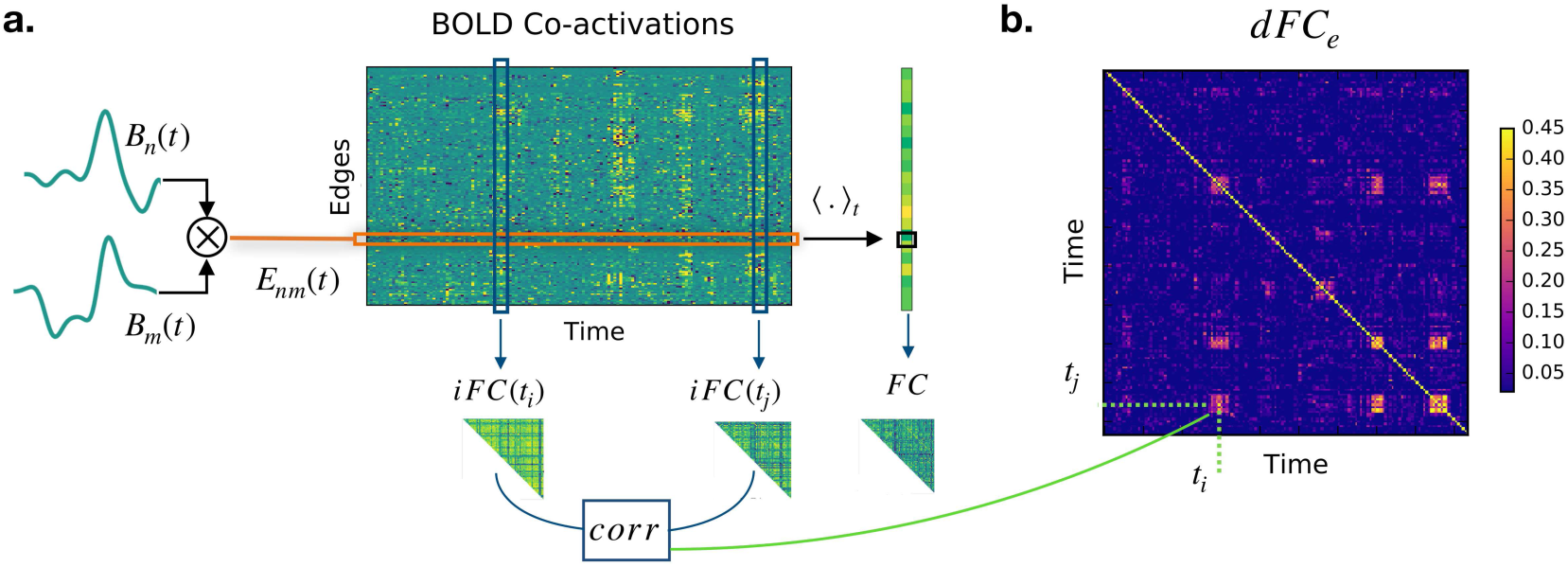
Two regimes of dynamic Functional Connectivity. (a) Given two nodes *n* and *m* the edge co-activation (CA) signal *E_nm_*(*t*) (orange box) is defined as the product of the z-scored BOLD signal *B_n_*(*t*) and *B_m_*(*t*). Averaging the BOLD-CA matrix over time we obtain the Pearson correlation across each pair of brain regions *n* and *m* (in black box, right), defining the static Functional Connectivity (*FC*). Each column of the BOLD-CA matrix represents and instantaneous realization of the Functional Connectivity (*iFC*). (b) The elements (*t_i_, t_j_*) of the edge-dynamic Functional Connectivity (*dFC_e_*) matrix are defined as the Pearson Correlation between *iFC*(*t_i_*) and *iFC*(*t_j_*). Note in panel (a) the presence of transient bouts of strong BOLD-CA (e.g., in the blue boxes). During these events, the *iFC* remains relatively correlated for few consecutive time points, which gives rise to diagonal (yellow) blocks in the *dFC_e_* matrix. The same CA burst (e.g., at *t_i_*) can re-occur in time after long periods of time (e.g., at *t_j_*), which gives rise to an off-diagonal *dFC_e_* block (e.g., at the crossing of the dashed lines in panel b).

In fact, averaging the edge CAs across time defines the Pearson correlation across each pair *n* and *m* (black box in Figure 2a, right). The correlation for each couple of regions (i.e., each edge) defines the static *FC* (Figure 2a, right). Thus, we can interpret the columns in Figure 2a (blue boxes) as instantaneous realizations of the Functional Connectivity (*iFC*) at different times *t*. The edge dynamic functional connectivity (*dFC_e_*, Figure 2b) is defined (without the use of sliding windows) by the Pearson Correlation of the *iFC*s at each couple of times *t_i_* and *t_j_*. On the one hand, the window approach to *dFC* allows a more reliable measure of the correlations to the detriment of the resolution over the temporal structure. On the other hand, the edge approach is more sensitive to spurious correlations but, crucially for our following analysis, it maintains the full-time resolution of the BOLD signals. Notably, the example Figure 2a extracted from a regime of interest, reveals that short duration bouts of strong BOLD co-activations (vertical stripes) spontaneously appear in association with a non-trivial *dFC_e_*. As a final remark, we also notice that the *dFC_e_* blocks (Figure 2b) occur in coincidence with the strongest edge co-activations, an observation that will have important consequences in our final analysis.

### 3.3 Multiple pathways to simulate RSN dynamics

To search for regimes of dynamic resting-state activity, we explore the parameter space by varying the global synaptic coupling *G* and the noise *N*, representing the impact of the structure over the local dynamics, and the stochastic currents simulating a generic environmental noise (thermodynamic, chemical, synaptic…), respectively. Then, we look for regions of the parameter space where a non-trivial functional network dynamics emerge. We define the switching index i.e., the variance across all elements of the upper triangular part of the *dFC_w_* matrix (Figure 3) to quantify the temporal irregularity of functional activity (roughly speaking, the number of different *dFC_w_* block structures and how often they switch between one another). We identify two regimes giving rise to *RSN* dynamics, where the respective neuroelectric correlates are qualitatively different.

- In the first case, a weak structural coupling (low *G*) is insufficient to promote any region into a high firing rate state, because the excitatory drive received by any node is relatively small. However, all regions can transiently reach a high firing rate state because the stochastic drive is strong. We refer to this as the *monostable* regime (Figure 3, bottom left).
- In the second case, the stronger structural coupling pushes a subset of regions into a strongly active state leaving the remaining regions in the down state. A weak stochastic drive ensures stable dynamics with few regions jumping between the up and down states. We call this the *bistable* regime (Figure 3, bottom right).

**Figure 3:**
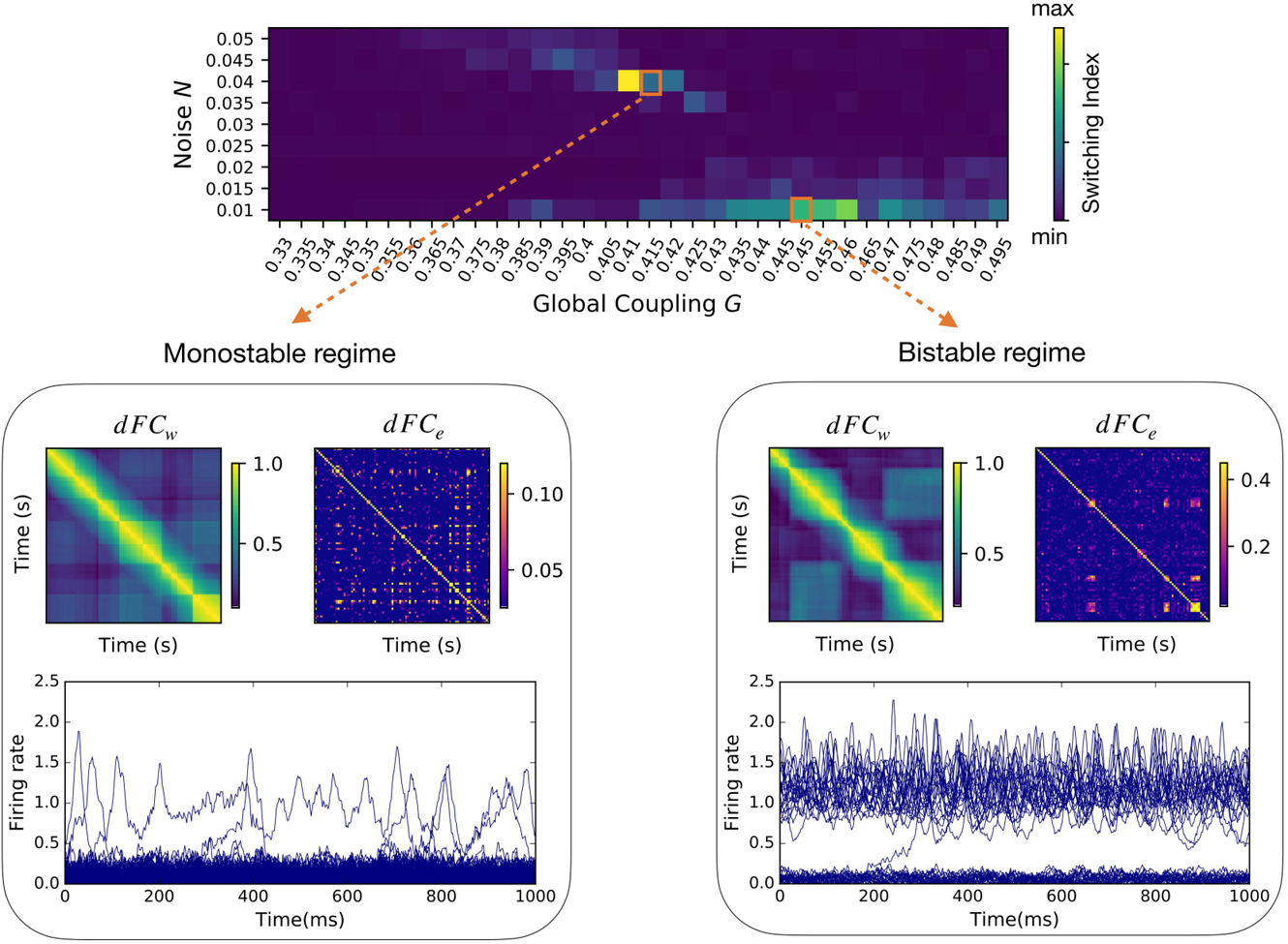
Two qualitatively distinct regimes of non-trivial functional dynamics. For every couple of global parameters (*G, N*) we calculated the dynamic functional connectivity (dFC) in a sliding window approach (*dFC_w_*; as in Methods) and in an edge-centric approach (*dFC_e_*; as in Figure 2a-b). The ‘switching index’ of each *dFC_w_* matrix, was evaluated as the variance of the respective upper triangular elements. We find two regimes of activity, named *monostable* and *bistable*, where qualitatively distinct neuroelectric organizations give rise to large-scale functional dynamics characterized by a non-vanishing switching index. In both regimes, the *dFC_w_* and *dFC_e_* display off-diagonal blocks, demonstrating a correlation between the functional activity at distinct times. The low global coupling *G* in the monostable regime (bottom left) does not guarantee a strong communication between the brain network regions, which most of the time populate the low firing rate (‘down’) state. A strong noise *N* pushes the brain regions in the high firing rate (up) state for short transients. A higher value of the global coupling in the bistable regime (bottom right) promotes a subgroup of regions in the high firing rate (‘up’) state. Low levels of noise perturb the equilibrium of the system provoking localized switching in both up→down and down→up directions (e.g., at *t* = 200ms).

In conclusion, the balance between the local dynamics and the global connectivity (tuned by *G*), together with appropriate levels of perturbation of the system (*N*), allow the same NMM model to generate diverse large-scale organizations qualitatively similar to those measured empirically.

Before moving on, let us notice that the block structures in the *dFC_w_* (examples in Figure 3, bottom panels) give the impression that whole-brain dynamics is organized by a sequence of transient but long-lasting stable periods of correlated activity. However, the analysis of the corresponding *dFC_e_* reveals that whole-brain dynamics is in fact characterized by much smaller periods of co-activations. Since we want to compare resting-state dynamics to the underlying fast neuronal activity, we are interested in the highest temporal resolution available. Therefore, unless differently specified, in the following analysis we use *dFC_e_*. We also notice that when *dFC_w_* is used, as the window slides in time, it will capture the transient edge co-activation bursts as long as the latter is present in the window, giving rise to spurious long-lasting block structures (e.g., see the different block organization between *dFC_w_* and *dFC_e_* in Figure 3, bottom right). Next, we analyze the neuronal mechanisms underlying resting-state functional dynamics generated by the model.

### 3.4 A generative mechanism for slow cascades of neuronal activations

Functional dynamics and the underlying neuronal activity fundamentally differ in their intrinsic time scales. Neuronal fluctuations typically evolve at the milliseconds scale. RSN dynamics unfolds in the order of tens of seconds. In this section, studying the results of our simulation, we reveal a potential mechanism by which local neuronal perturbations can generate a slow cascade of activation across the brain network, thus approaching the temporal scale of RSN dynamics.

To do so, we introduce a framework to interpret the simulated firing rate dynamics and its dependence on the structural connectome. We focus on the bistable regime where the low noise level allows a clearer visualization of the dynamic mechanisms in act. The hypotheses developed here will be later tested on the monostable regime as well as in empiric data. We identify five main categories of nodes based on their simulated firing rate activity:

- (D) *down* regions (in light blue in Figure 4a) display a low firing rate state throughout the duration of the simulation, as the up state is practically unreachable by noise-driven fluctuations. According to Section 3.1, the separatrix lies close to the up state (Figure 4,b, top). Their low firing rate regime (Figure 4,b, bottom) makes them poor communicators in the network hierarchy.
- (U) *up* regions (in light red in Figure 4a) always show a high firing rate (Figure 4,b, bottom), constantly providing inputs to their targets. The separatrix is close to the down state and is never reached (Figure 4,b, top).
- (J) *jumping* regions (in green in Figure 4a) undergo regular local transitions, dwelling for a relatively long time in both low- and high-firing rate states (Figure 4,b, bottom). In the projected phase space of these regions, the separatrix lies midway between the up and down states (Figure 4,b, top). Therefore, they have the same probability to jump from up to down and vice versa Note that the timing of the (J) jumps defines a new *slow* time scale for the system. The fact that these jumps occur on a regular basis during the simulation ensures that this time scale is ever-present in the large scale network dynamics.
- (D*) *down-up* regions (in dark blue in network plot Figure 4a) have a stable activity around the down state fixed point but–in rare occasions–manage to reach the up state stable focus for a certain transient time.
- (U*) *up-down* regions (in dark red in network plot Figure 4a) have a stable activity around the up state but are occasionally driven into a short-lived excursion to the down state.

**Figure 4:**
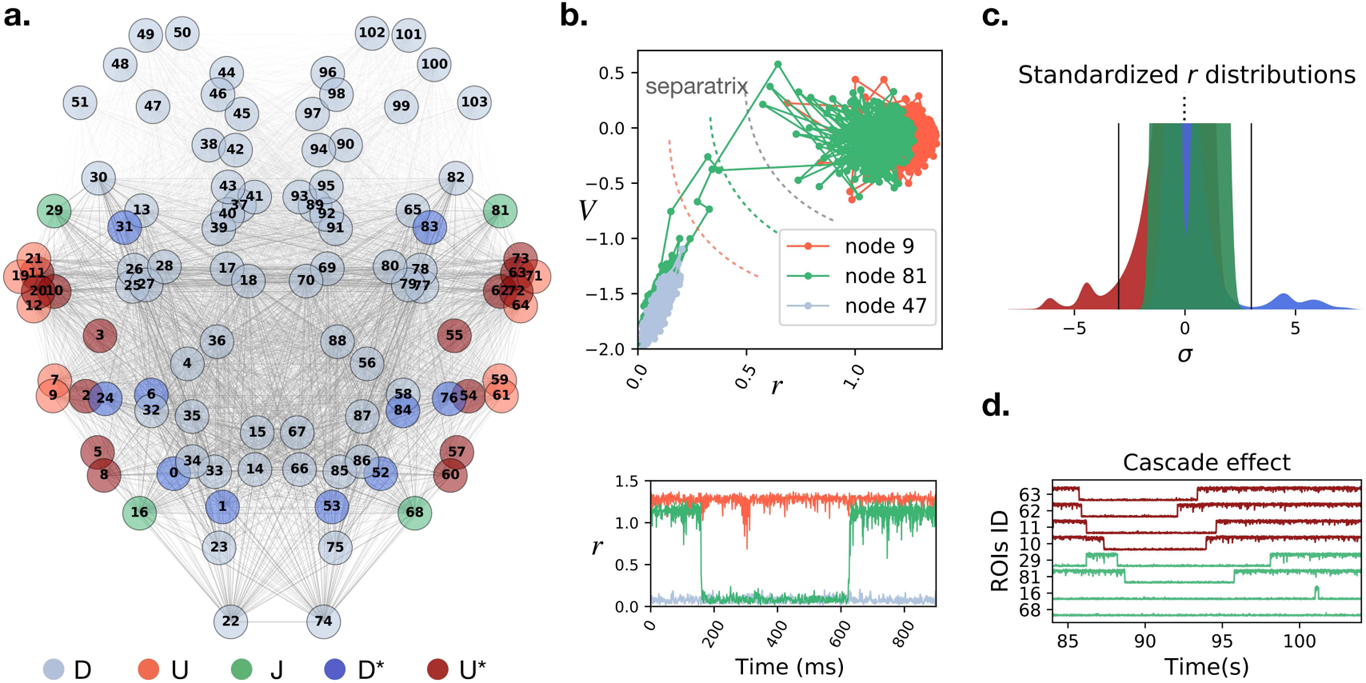
Mechanisms of cascade generation in the synthetic model. (a) Different regions have a different fate depending on their location in the connectome. We classified the regions in five classes (D, U, J, D*, U*) according to their activity. (b) Example exploration of the projected 2D phase space (top) and firing rates activity (bottom) of the ‘up-U’ (light red), ‘down-D’ (light blue) and ‘jumping-J’ (green) regions. (c) Distribution of the standardized firing rates in different classes. Class (J) regions have two modes but never cross the ±3*σ* threshold (black lines). Class (U*) (dark blue) and class (D*) (dark red) regions dwell most of the time in the up and down states, respectively. Only in important rare occasions the *-regions cross the threshold to jump on the other side, substantially deviating from their baseline activity. The leading role of the *-regions as compared to the other classes is shown using Principal Component Analysis (PCA) in Figure 5-1a-b. (d) Example of a cascade: when the (U*) node 63 jumps into the down state, it first drags down the node 62 (with which it shares the strongest structural link in the network). After them, other strongly connected nodes follow the trend.

The topographic organization of the firing rate classes shows the influence of the connectome in organizing whole-brain dynamics (Figure 4a; regions labels in Table 1). Nodes receive different inputs as a function of the location in the connectome, which grants a different jumping probability and therefore a different role in network dynamics. A stochastic jump is a functionally important event for the network, as it corresponds to a sudden firing rate increase or decrease, which greatly influences the activity of downstream nodes. Thus, since the nodes of class (J) can jump regularly, we expect them to have a major role on whole-brain dynamics. A Principal Component Analysis (PCA) of the regions’ firing rates shows that nodes (J) are the main contributors to the first three principal components, explaining most of the system variance (explained variance ratio*>* 0.59; see Figure 5-1a).

**Figure 5:**
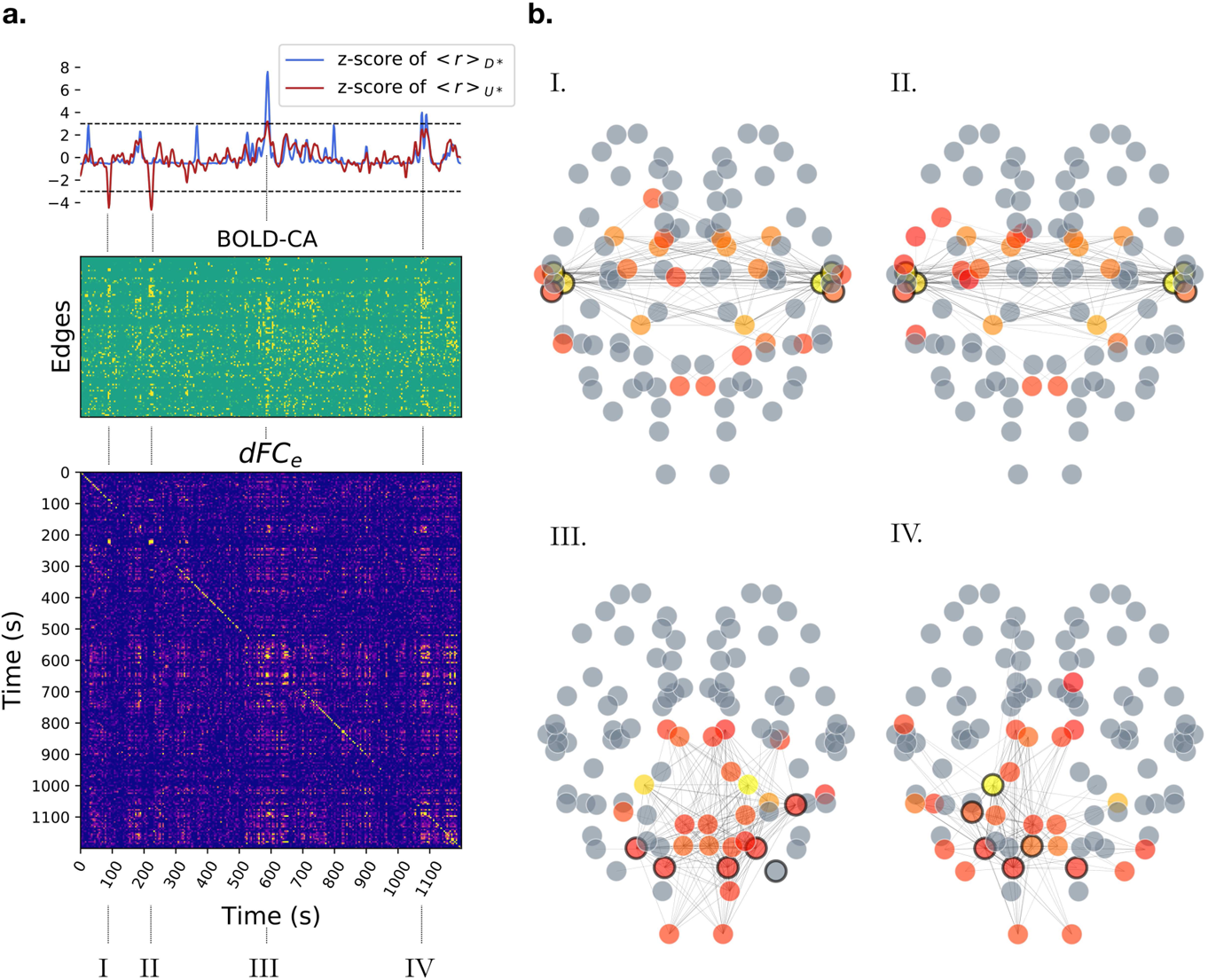
Resting-State Network formation. (a, top) The standardized firing rate activity in the (U*) and (D*) classes (class-specific average; dark red and dark blue, respectively) is characterized by peaks (the strongest are marked as I, II, III, IV) occurring in correspondence of cascades similar to Figure 4d. (a, middle) During a cascade we also observe a peak of BOLD co-activations (CA), appearing as vertical strips. Many, but not all, edges are recruited. (a, bottom) The blocks in the *dFC_e_* matrix appear in correspondence of CA events, showing that these bursts generate stable epochs of functional connectivity correlated in time. (b) In each selected epoch (I, II and III, IV), the large firing rate cascades trigger the jump of other nodes away from baseline activity (circled in black) and promote specific functional hubs at the BOLD level, represented by colored nodes in the network plots. A functional hub is defined by the components of the first leading eigenvector (linear combination of brain regions explaining most of the variance in the data; eigenvalue *λ >* 0.41) associated to the istantaneous *iFC*s at times *t_I_, t_II_, t_III_* and *t_IV_*, respectively. The most representative hub regions are depicted in yellow. Gray regions have been excluded as they do not contribute substantially. Only the edges with the highest co-activations are displayed. Importantly, CA events generated from neuronal cascades at specific sites support distinct functional networks which are not correlated among themselves (e.g., no off-diagonal *dFC_e_* block between I and III).

Another key element of the simulated network dynamics is the occurrence of rare events, identified by large deviations e.g., above three standard deviations *σ* away from baseline firing rate (black lines in Figure 4,c). Given their regular jumping, the activity of regions (J) never grows above 3*σ* (bimodal distribution of standardized firing rate in green in Figure 4,c). The same holds for (D) and (U) regions. In contrast, we find that the regions in classes (D*) and (U*), occasionally grow above 3*σ* during their rare jumps across the separatrix (Figure 4,c, dark blue and dark red distribution, respectively). Our simulation shows that these rare events can trigger cascades of transitions across the separatrix of a subset of regions (e.g., Figure 4,d). The propagation of a cascade depends on the structural location of the source node. Let us consider the case of a node with jumping potential, which is also a hub for the network (i.e., with high centrality; (van den Heuvel & Sporns, 2013)). Its local reshaping will have a wide influence throughout the network. However, the hubness of a jumping node does not guarantee the crossing of the separatrix by its target regions. If the hub is connected to a very stable subset of regions, following a jump of the hub, most of the separatrix lines in the targets will be slightly shifted, but not enough to allow any crossing. In contrast, if there exists a subset of strongly connected nodes with jumping potential, the jump of one will have strong effects within such subnetwork. In order to illustrate this point, let us consider that a group of nodes from class (U*), have very strong links binding them. Then, the occasional jump down of one of these nodes will strongly shift the separatrix of the other (U*) nodes towards the up stable focus, increasing their probability of jumping down. Then, many (U*) nodes will be dragged down one after the other, facilitating the next jumps, producing a cascade effect (as in Figure 4d). During this event, the (J) nodes get also involved, interrupting for some time their regular control over the network. A windowed analysis of the first principal component of the firing rate activity shows that the nodes in the *★*-classes are driving the system during these rare events (Figure 5-1b).

The cascade effects quickly drag the system away from its standard state, establishing short epochs of increased deviations from baseline activity (the average firing rate in classes D* and/or U* deviates from baseline; Figure 5,a, top). In our simulation we mark four of these epochs (I, II, III, IV; Figure 5,a, bottom; Figure 4d refers to epoch I) which are separated by long periods of standard activity.

Summarizing, we have described how the jumps of the firing rate at local sites are extremely relevant for the unfolding of the microscopic activity throughout the brain, as they provoke the largest perturbations in the system. If jumps happen with a certain regularity, they will generate a slow time scale in the system. This slow rhythm in our simulations can be traced back to a small subset of regions (J) which transit between the up and the down states. These regions lead the baseline evolution of the entire system, acting as homeostatic agents which keep the system dynamics stable. On rare but important occasions this control weakens, which allows the propagation of a large cascade of activity. Such cascade effect takes place when a sudden local transition brings a typically stable region ((U*) or (D*)) away from baseline activity and induces other regions to reorganize. The cascade goes on until the noise and the structural pressure take the system back to its normal evolution e.g., through the normalizing action of the jumping regions (J). We then explore whether such rare deviations from baseline activity account for resting-state brain dynamics.

### 3.5 Neuronal cascades activate distinct RSN

The previous analysis focused on the simulated (high time resolution) neuronal activity. We now look at the simulated (low time resolution) BOLD, which signals are characterized by collective co-activations (CA) (Figure 5a, middle). Interestingly, these events are aligned to the large deviations from baseline of the firing rate activity of the (U*) class or the (D*) regions (Figure 5a, top, segments I,II and III, IV, respectively). When looking at the corresponding *dFC_e_* matrix (Figure 5a, bottom), we also note that the most stable *dFC_e_* blocks correspond to the largest BOLD-CA events. This observation is not a triviality, since a higher co-activation does not imply a stronger correlation. The presence of off-diagonal blocks in the *dFC_e_* shows that CA patterns are correlated when generated either by the (U*) or in the (D*) regions. In this example, the CA patterns generated by the jumps of distinct classes are not correlated. Each CA pattern defines the upper triangular part of an instantaneous functional connectivity *iFC* (as in Figure 2a). Figure 5b displays, for each selected epoch, the edges with the highest CA values. The functional hubs, a subset of regions with a central role in the functional dynamics, are defined in each epoch by the *iFC*’s leading eigenvector (Melozzi et al., 2017), and represented as different colors in the brain network plots (from red to yellow in a scale of importance). We highlight with a black rim the nodes whose firing rate deviated from baseline activity above a threshold (fixed at 3*σ*). Notice that while the cascade generates at certain network locations (black rims), the dynamics of the RSN establishes a new emergent phenomenon at the large scale (colored nodes). The same perturbation can re-occur spontaneously eliciting similar functional networks correlated at distinct times, as it is marked by the off-diagonal blocks of the *dFC_e_* (network plots I, II or III, IV). Summarizing, neuronal perturbations starting at different locations in the connectome elicit different BOLD-CA events, which result in distinct functional networks.

### 3.6 Neuronal cascades and BOLD co-activations in empiric data

In the previous sections, we described how the large spontaneous deviations from baseline activity play a key role in the simulated system dynamics by activating specific large-scale functional networks. Here we hypothesize that a qualitatively similar network behavior takes place in empiric resting-state data (see Discussion). To test this, we provide a working definition of *neuronal cascades* as a global measure of such long-lasting perturbations of the neuroelectric activity. See Discussion). For illustrating the pipeline, we use an example trial from an EEG/fMRI resting-state human dataset (See Methods). First, we binarize the firing rate activity in every brain region by assigning a unitary value when its activity exceeds ±3*σ* (Figure 6a). Then, we define the magnitude of the deviations from baseline by summing the binarized EEG over the regions (gray line in Figure 6b, top). Finally, we convolve the obtained signal with a Gaussian kernel (width = 1.94*s* = BOLD Time Repetition) and we downsample it to the same time resolution of the BOLD signals to allow comparison. The resulting neuronal cascade signal (blue line in Figure 6b, top) is characterized by bumps which describe a long-lasting increase in the magnitude of non-standard perturbations (e.g., above 3*σ*). Neuronal cascades should not be confused with the concept of *neuronal avalanches* (Beggs & Plenz, 2003), which consist of consecutive deviations from baseline activity of a subset of brain regions or localized neuronal groups (example of a neuronal avalanche in the red box, Figure 6a, bottom). The number of regions in the subset defines the size of an avalanche, while the duration of the consecutive activations defines its lifetime (up to few hundreds of milliseconds in empiric data). Neuronal cascades are a global measure that describes a slower process on the order of tens of seconds. In Figure 6b (bottom panel) each red dot marks the occurrence of an avalanche of a certain size (dot size represents the duration). Remarkably, neuronal avalanches accumulate in correspondence with the largest neuronal cascades. Thus, we can think of the neuronal cascades as the clustering of neuronal avalanches. This first major result proves that the probability of observing strong brain fluctuations (including neuronal avalanches) is not constant in time, but increases during neuronal cascade peaks, which occur at an infra-slow time scale. Also, our simulation shows localized increases in cascade activity, which are shorter in the noisy monostable regime and more prolonged in the bistable regime (Figure 7a).

**Figure 6:**
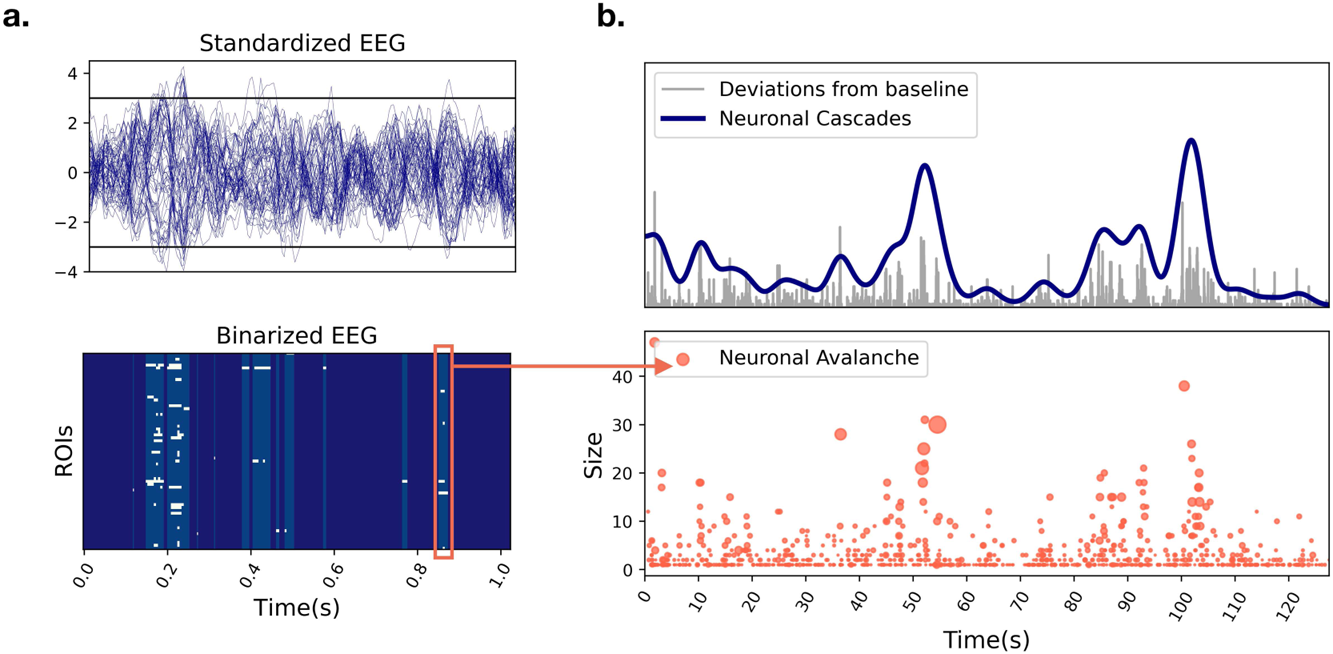
Neuronal cascades and neuronal avalanches. (a) Standardized EEG activity extracted from a resting-state human EEG/fMRI dataset (top). The activity is binarized assigning a unitary/null value every time the activity in a region is above/below a certain threshold (e.g., ±3*σ*; black lines). The obtained binary raster plot (bottom) is characterized by intermittent epochs of deviations from baseline activity. Neuronal avalanches are defined as consecutive deviations from baseline activity (e.g., red box). (b) We extract the global magnitude of the deviations from baseline (top, gray signal) by summing the binary EEG raster plot over the ROIs. This signal is convoluted with a Gaussian kernel (width = 1 BOLD Time Repetition (TR)) and downsampled to obtain the same resolution of the BOLD activity, which defines the neuronal cascades signal (blue). Neuronal cascades can be thought of as clustering of high magnitude avalanches, whose occurrence in time is not homogeneous (bottom).

**Figure 7:**
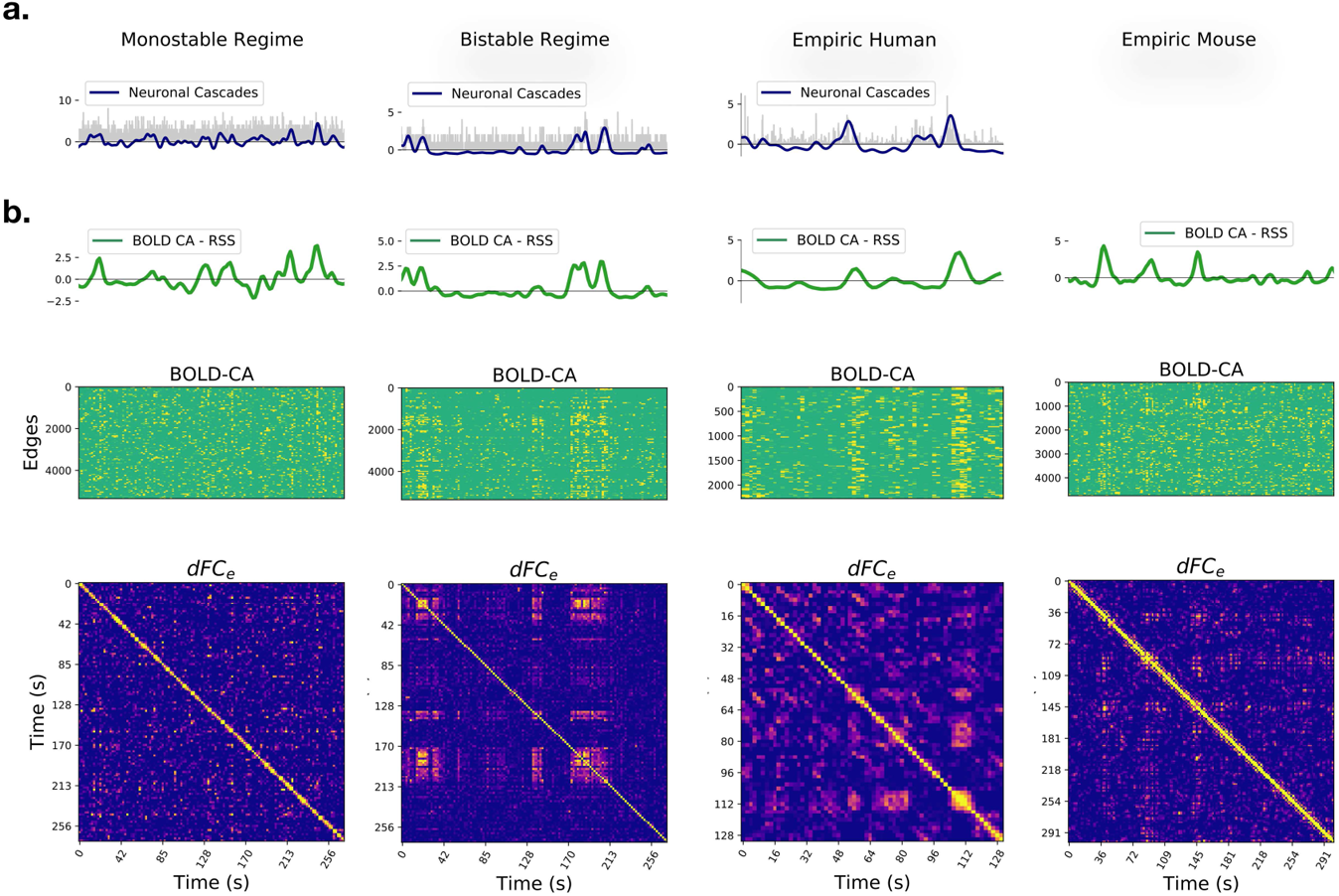
Neuronal cascades drive the functional dynamics. (a) Example of neuronal cascades in the monostable and bistable synthetic regime and for a representative subject of the empiric EEG/fMRI Human dataset. (b) The BOLD-CA in simulated and empiric mouse and human datasets (middle panels) are characterized by sudden collective events involving large network parts (vertical stripes). The root sum square of BOLD-CA across all edges (RSS, green lines, top panels) defines the global CA amplitude signal for each dataset. Concurrently, the *dFC_e_* matrices (bottom panels) display both diagonal and off-diagonal blocks, remarking the non-trivial re-occurrence of the same stable functional network at distinct times (see Figure 2b). At a visual inspection, the BOLD-CA events happen in coincidence with *dFC_e_* blocks and, most notably, the neuronal cascades and RSS signals (blue and green lines in panels (b-c), respectively) co-fluctuate in most instances.

At the level of fMRI, another *in-silico* prediction is the presence of strong BOLD-CA events and their co-occurrence in coincidence with off-diagonal *dFC_e_* blocks. We verify the presence of BOLD-CA events in synthetic, empiric mouse fMRI and in human EEG/fMRI datasets (Figure 7b, central panels), suggesting that temporally inhomogeneous bursts constitute a generic dynamic modality of real(istic) brain networks. To show the relation with *dFC_e_*, we first extract from the BOLD-CA signals the root sum squared (RSS) time series, which quantifies the amplitude of all edge co-activations at each time point (Figure 7b, green lines, top panels). Then, we partition the RSS signal in CA *events* and nCA *non-events* (respectively above and below the 98th percentile of the RSS values). We show that the correlation between any two events is statistically higher than the correlation between events and non-events as well as between couples of non-events (Figure 8a). The results hold for synthetic and empiric results, which were pooled over several experimental trials. Since the correlation between co-activation patterns at different times correspond to the off-diagonal elements of the *dFC_e_* matrix (as shown in Figure 2a-b), we conclude that bursts of BOLD-CA events account for the highest off-diagonal values in the *dFC_e_* in experimental datasets, as predicted by the model. In Figure 8b we show an example of empiric human *dFC_e_* (top), and the same matrix where times are sorted accordingly to increasing RSS. Green lines represent different percentiles (50th, 75th and 95th) of the RSS. Notice that, in line with the results above, the non-trivial temporal correlations (yellow off-diagonal *dFC_e_* elements) involve those time frames with the highest network co-activations.

**Figure 8:**
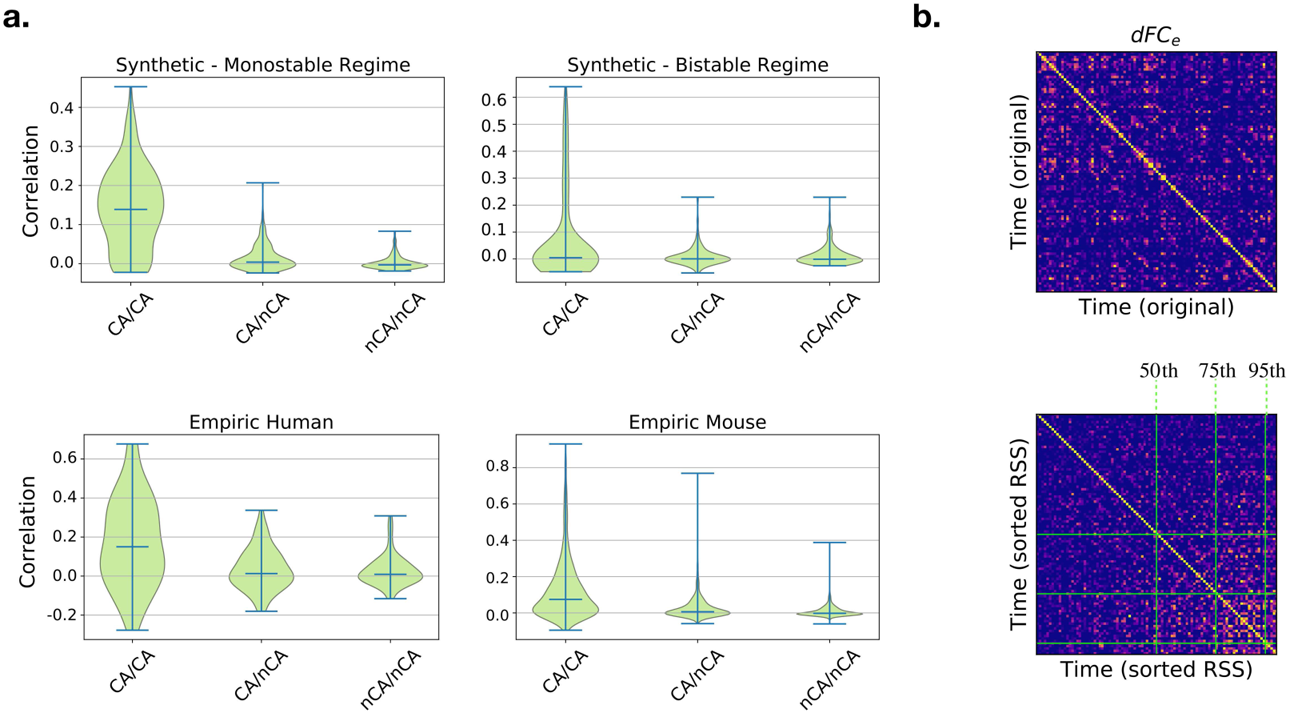
(a) The largest BOLD co-activations events (CA, above the 98th percentile of the RSS, green line in Figure 4c) are distinguished from *non-events* (nCA, below threshold). We report the synthetic and empiric correlations between *iFC*s at times within CA events (left in every panel), between CA events and non-events nCA (center of panels), within non-events nCA (right of panels). These correlations are by definition the off-diagonal values of the *dFC_e_* matrix (see Figure 2a-b). The distribution of the correlations within events is wider and explains the greatest off-diagonal correlation values of the *dFC_e_* across all the synthetic and empiric datasets. This principle is explicitly shown in panel (b), where the original *dFC_e_* extracted from an empiric human trial (top) was sorted according to increasing RSS (bottom), leading to the clustering of high correlations towards high CA times. This shows that most of the non-trivial temporal correlations involve co-activation times falling in the last quartile of the RSS (above the 75th percentile, central green line). Thus, the strongest co-activation events drive the dynamics of *FC*.

### 3.7 Neuronal cascades subtend RSN dynamics

Based on the theoretical results of the previous sections (summarized in Figure 5), neuronal cascades should give rise to BOLD-CAs. In fact, a visual inspection of simulated and empiric data suggests the co-occurrence of neuronal cascades (blue lines in Figure 7a) and BOLD-CA (RSS; green lines in Figure 7b). Remarkably, a similar correspondence is present in an EEG/fMRI resting-state human dataset (best trial shown in Figure 7), all the more since the simulations are done using a mouse connectome. To characterize this correspondence, we correlate the cascades signal with the BOLD-CAs amplitude profile. In both the monostable and the bistable regimes, the two measures are significantly correlated (*ρ* ∼ 0.54 and *ρ* ∼ 0.91, respectively). In Figure 9a (top) we report the correlations between the cascades and the BOLD-CA amplitudes across several trials of a human cohort (see Methods section for trial selection). The correlation is carried out by shifting the RSS time series of a given time-lag. A peak of correlation appears naturally between the two measures when the BOLD signal is shifted backwards by 2 time points (3.88*s*) (Figure 9a, bottom; blue line). Therefore, as expected, the EEG neuronal cascade signal precedes the BOLD co-activations by a few seconds. Notice that, in general, a shift forward of the BOLD activity results in a rapid loss of correlation.

**Figure 9:**
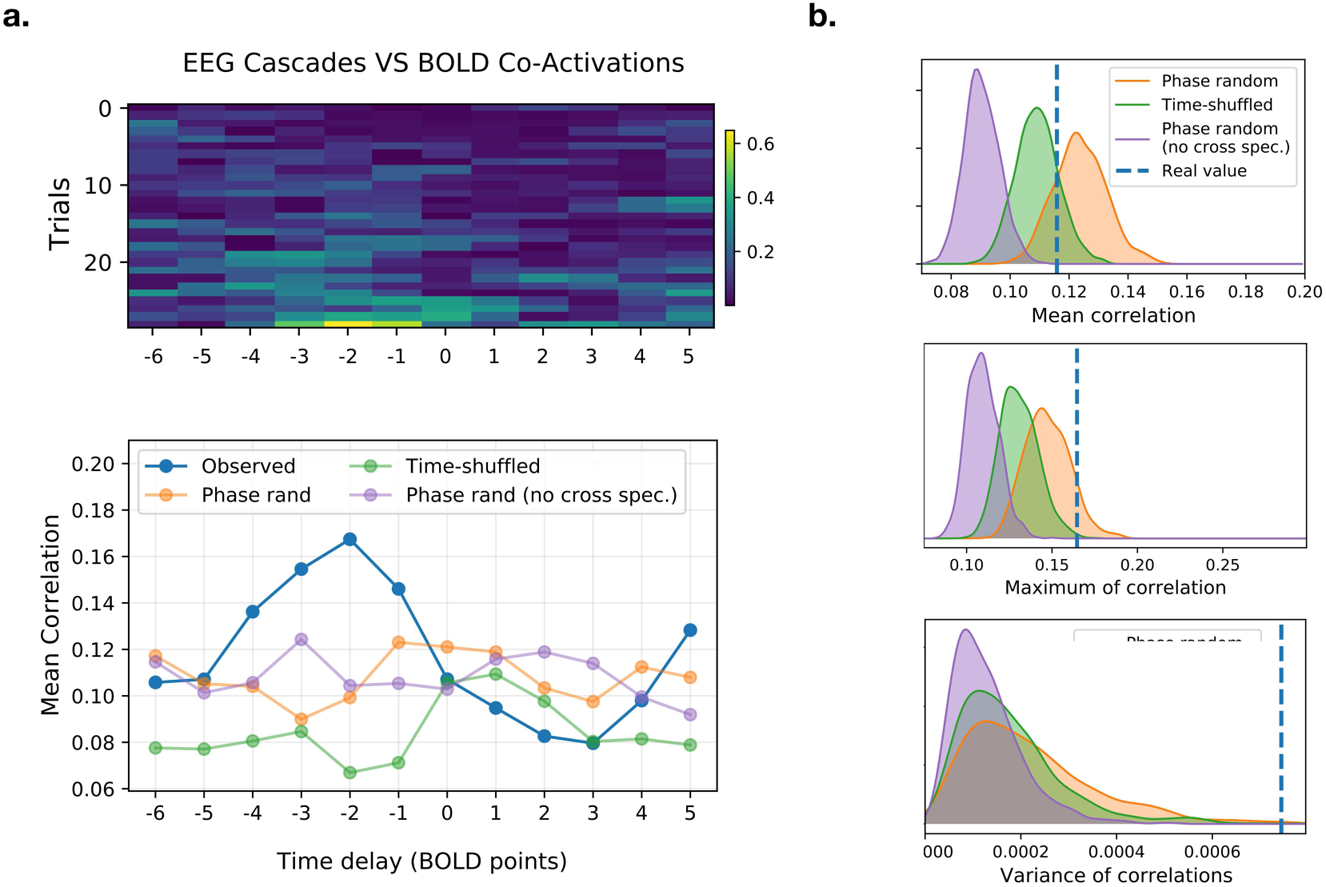
(a, top) Correlation between the cascade magnitude and the BOLD-CA amplitude (blue and green lines in Figure 4b-c) for different time lags in several EEG/fMRI trials extracted from a Human cohort of resting subjects. Negative lag is associated with a shift backward of the BOLD signal. (a,bottom) The cross-correlation averaged across trials shows a clear trend (blue line). The peak of correlation at lag −2 sampling points (1*pt* = 1.94*s*), as well as the rapid fall for positive lags, confirms that the EEG precedes the BOLD activity by few seconds. The same profile is evaluated by comparing the largest cascades with the BOLD-CA signals extracted from 1000 time-shuffled (example in green), 1000 phase-randomized (cross-spectrum preserved, example in orange), and 1000 phase-randomized (cross-spectrum not preserved, example in red) BOLD surrogates for every subject (see Figure 9-1a-b for surrogate properties). (b) The distribution of the mean, maximum, and variance of the cross-correlations for each surrogate model is displayed and compared to the empiric values. In particular, the variance plot shows a clear significance of the results (single subject results in Figure 9-1c).

Finally, to prove the statistical significance of our observations, we build surrogate BOLD signals and we compare the associated CA amplitudes to the neuronal cascade signal in each trial. In particular, we compare the observed cross-correlation profile (blue line Figure 9a, bottom) with the profiles extracted by correlation of neuronal cascades and surrogate CA amplitudes of three kinds:

- Time-shuffled: Example of cross-correlation profile in Figure 9a, green line.
- Phase-randomized, cross-spectrum preserved: Example in Figure 9a, orange line.
- Phase-randomized, cross-spectrum not preserved: Example in Figure 9a, purple line.

In all the surrogates, the functional dynamics is disrupted (see Methods). In the first two models, the static FC is preserved but its dynamic evolution is made trivial. Namely, the time-shuffled model consists of random *FC* jumps around a fixed pattern, while in the phase surrogate (cross-spectrum preserved), stationarity is strictly imposed by destroying any coherent fluctuation around the static *FC*. In fact, the latter corresponds to preserving the static correlations across node pairs i.e., the average of the edge co-activation signals. These two surrogates also preserve the original fat-tail distribution of the BOLD CA amplitudes (RSS values), and therefore supports bursty CA events (Figure 9-1a-b). When the cross-spectrum is not preserved, the static FC is corrupted and the CA amplitude distribution is normalized. For each surrogate type, we compute 1000 cross-correlation profiles. The distribution of their mean, maximum, and variance (averaged across all the trials) is shown in Figure 9b. The observed values are always significant when compared with the Phase-randomized surrogate with cross-spectrum not preserved. The observed mean values lie between the random (time-shuffled) and the ordered (phase-randomized) scenario (Battaglia et al., 2020). Only the Phase-randomized surrogate, where the cross-spectrum is preserved, reaches the peaks of correlation of the observed data, and the significance of the observed maximum is *p* = 0.108. The variance of the correlation profile is always significant (*p* < .001) as compared to all the surrogate models. The result holds in several instances at the single-subject level (Figure 9-1c). We conclude that neuronal cascades drive BOLD-CAs dynamics thus shaping whole brain resting-state functional connectivity.

## 4 Discussion

Cognitive function requires the coordination of neural activity across many scales, from neurons and circuits to large-scale networks (see Betzel and Bassett (2017) and references within). As such, it is unlikely that an explanatory framework focused upon any single scale will yield a comprehensive theory of brain activity, cognitive function or brain disease. Fast time scales evolve within the quasi-stationary constraints imposed by slow time scales. As the latter changes, the organization of the former must change consequently. Similarly, microscopic neuronal events can trigger large-scale complex responses (Houweling & Brecht, 2008; Huber et al., 2008). We demonstrated this principle along the organization of the fast neuroelectric correlates of the slowly evolving large-scale dynamic Functional Connectivity (*dFC*) patterns for two mammalian species. Large neuronal cascades, defined by collective deviations of the simulated EEG signals away from baseline activity, can emerge as BOLD co-activation (CA) events, i.e. large simultaneous activation at pairs of regions across the brain network. In turn, the intermittent set of BOLD-CA events underlies specific stable *FC* epochs. Thus, we suggest that large neuronal cascades have an organizational role in the resting-state brain dynamics.

In this work, we have adopted a theory-driven approach to reproduce qualitatively the resting state *FC*, analyze its dynamics and trace it back to its neuronal correlates. The neural mass model used to simulate regional dynamics maps the activity of an infinite number of all-to-all coupled neurons, each described by a simple phase-oscillator equation, into the mean-field firing rate and membrane potential variables (Montbrío et al., 2015). Although not giving full access to biophysical processes, as when using detailed single neuron models, the model we used preserves important information about the microscopic neural network organization of a simulated brain region (e.g., the neuronal synchrony; see Methods). The low computational load enables the simulation of fMRI-like sessions in TVB. Future applications allow the possibility of including synaptic dynamics (Taher, Torcini, & Olmi, 2020), adaptation (Gast, Schmidt, & Knösche, 2020), excitatory versus inhibitory populations (Dumont & Gutkin, 2019; Laing, 2017; Montbrío et al., 2015), among others. The use of these models establishes a new venue to address multiscale phenomena, maintaining the co-existence of microscopic and macroscopic scales of organization (Coombes & Byrne, 2019).

To simulate the spontaneous emergence of dynamic functional networks we tuned each neuronal population into a bistable regime. We expect similar results to hold for other models supporting local bistability. In our case, we can directly interpret the two stable states as two configurations of the spiking neural networks. Varying the global couplings, we discovered two regions of the parameter space–defining the monostable and the bistable regimes–where the large-scale organization is dynamical, but qualitatively different. In both regimes, the neuronal activity is characterized by up and down states, which occurrence was observed both in vitro (Cossart, Aronov, & Yuste, 2003; Plenz & Kitai, 1998) and in vivo under several conditions such as anaesthesia, slow-wave sleep, quiet waking and also during perceptual task across several species (Engel et al., 2016; Jercog et al., 2017; Luczak, Barthó, Marguet, Buzśaki, & Harris, 2007; Steriade, Nunez, & Amzica, 1993; Vyazovskiy et al., 2011). The functional dynamics in the two regimes also differs e.g., in terms of the statistical properties of the BOLD-CA events (Figure 8a) or in the life-time of the stable *FC* states (i.e., the size of the *dFC* blocks in Figure 7b, bottom).

The generation of a variety of dynamic regimes from changes in the global ‘environmental’ parameters is in keeping with the degeneracy principle, according to which multiple models and/or parameter settings capture the neuronal variability (Marder & Taylor, 2011), and it is expected also in biological systems. For example, sleep deprivation, which alters brain and body function (Medic, Wille, & Hemels, 2017), is associated with changes in the speed of the *FC* evolution (Lombardo et al., 2020). Accordingly, distinct features of the *dFC* should correspond to a different underlying neuroelectric organization (and perhaps to distinct mechanisms of communication across brain regions (Battaglia & Brovelli, 2020)). The transition between wakefulness and sleep brain states offers an example of sudden change of the whole-brain dynamics associated with the microscopic neuroelectric re-organization (Boly et al., 2012; El-Baba et al., 2019; Larson-Prior et al., 2009; Tagliazucchi, Crossley, Bullmore, & Laufs, 2016).

While the debate over the actual nature of functional connectivity dynamics is still ongoing (Hindriks et al., 2016; Laumann et al., 2017; Líegeois, Laumann, Snyder, Zhou, & Yeo, 2017), several evidences were provided in favor of the *dFC* as a legitimate measure of the functional evolution (Preti et al., 2017). Here, we found that a large part of the functional information is condensed in large BOLD-CA events, in keeping with previous results (Zamani Esfahlani et al., 2020) (see Figure 5a). The occurrence of the BOLD-CA events cannot be ascribed to the spectral properties of the BOLD signals nor to motion artifacts (Zamani Esfahlani et al., 2020), which supports a genuine phenomenon. Our results provide an *in silico* support to a burst-based system dynamics where correlated CA events subtending RSNs occur intermittently and are separated by long low-activity periods. These results are in accordance with previous works showing that the relevant information about the major RSNs is carried by the largest fluctuations of the fMRI blood oxygenation level-dependent (BOLD) signals (Allan et al., 2015; Cifre, Zarepour, Horovitz, Cannas, & Chialvo, 2017; Gutierrez-Barragan, Basson, Panzeri, & Gozzi, 2019; Liu & Duyn, 2013; Tagliazucchi, Balenzuela, Fraiman, & Chialvo, 2012). The spontaneous bursts of BOLD-CA highlight recurring sets of network edges, which subtend special RSNs.

How the system is organized around these preferential sets and what triggers the sudden co-fluctuations remain to be determined. Our analysis remarks the central role of the structural connectivity in this process, in line with a recent work suggesting that CA are shaped by structural modules (Pope, Fukushima, Betzel, & Sporns, 2021). Here, we suggest that large neuronal cascades generated by sudden neuronal re-organization in source regions may play a central role in driving the exploration of the connectome and in determining the large-scale functional connectivity dynamics.

During the BOLD-CA events, the evolution of the *FC* slows down, as shown by the high correlation between functional states at consecutive times (Figure 7b, bottom panels). An emergent stable *FC* elicited by a BOLD-CA event can persist for several seconds. The same state can re-appear after a few minutes in coincidence with another CA. In many dynamical systems, slowdowns and large-scale events are typically observed at the transition between different states (or phases) i.e., at the critical point (Scheffer et al., 2009). Critical dynamic systems are characterized by other typical properties, including the presence of fluctuations at all spatiotemporal scales (Cocchi, Gollo, Zalesky, & Breakspear, 2017). Historically, the hallmark of a critical organization of the brain activity came from the observation of neuronal *avalanches* whose size and duration fit a power-law (Beggs & Plenz, 2003) (for a ‘critical’ look at brain criticality see Beggs and Timme (2012)).

In order to compare the fast EEG and the slower BOLD signals, we focused on the slow unfolding of the EEG deviations from baseline activity. In other words, instead of focusing on the size and duration of the neuronal avalanches, we analyzed their collective occurrence at the resolution of seconds, which we refer to as neuronal cascades (Figure 6). We showed that the cascade magnitude is correlated to the BOLD co-activation amplitude (Figure 9a). In particular, in the empiric human dataset we observe a rapid drop of this correlation when the BOLD activity is shifted backward w.r.t. the EEG signal. This fact makes our result more robust since we expect a change of the BOLD activity to happen *after* a sustained neuroelectric activity.

Notice that in the empiric data set, not all the neuronal cascades give rise to BOLD CAs, and not all the BOLD CAs are preceded by large neuronal cascades. In general, we should expect a non-trivial interaction between these phenomena and other infraslow processes. For example, a growing corpus of evidence relates the slow functional and neuroelectric dynamics with other physiological and cognitive infraslow processes such as neuromodulation (Shine et al., 2016), visceral (heart and gut) inputs (Azzalini, Rebollo, & Tallon-Baudry, 2019), and cognitive performance (Monto, Palva, Voipio, & Palva, 2008).

In the cases in which neuronal cascades and BOLD CA events co-occur, we can hypothesize that the stable *FC* state evoked by a CA event is related to the specific structural channels in which the neuronal cascade spreads. In fact, we showed in the simulated data that the largest neuronal cascades underlie specific functional patterns (Figure 5), depending on the location of the onset. Also, the observation that the same cascade appears at regular intervals (see for example the three peaks at second ∼ 100, ∼ 200 and ∼ 300 in Figure 5a) suggests that the re-occurrence of the same stable patterns in time can be traced back to the increased probability of the same cascade to occur following a first large event, in the same way an after-shock follows the main earthquake. It is interesting to note that, as in our model, correlated bursty events associated with memory effects are observed in many natural systems (Karsai, Kaski, Barabási, & Kertész, 2012).

The above hypotheses require further exploration of larger empirical data sets. In the model, we have shown that the multiscale mechanistic origin of large cascades can be understood ultimately by the interplay between the local neuronal organization, the stochastic nature of its sudden re-organization and the structural constraints it obeys. The present work provides a new understanding of whole-brain functional dynamics, which is shaped by neuronal cascades giving rise to large BOLD co-activation. Since most neurological disorders are characterized by complex reorganizations at the neuronal scale, it will be interesting to determine whether specific alterations in neuronal cascades sign specific neurological disorders, and explain the already identified alterations in whole-brain dynamics.

## 5 Materials and Methods

### 5.1 Empirical Mice and Human data sets

The connectome used for the simulations was extracted using The Virtual Mouse Brain pipeline described in (Melozzi et al., 2017), which processes the tracer experiments performed at the Allen Institute (Oh et al., 2014). There, adult male C57Bl/6J mice are injected in specific brain regions with a recombinant adeno-associated virus, which expresses the EGFP anterograde tracer. In each experiment, the tracer migration signal is detected by a serial two-photon tomography system. The anterograde tracing provides information about the axonal projections starting at the injected site. We define the injection density of the source region as the number of infected pixels normalized by the total number of pixels in the region. Similarly, the projection density is defined in every region as the number of pixels detected in the target region following an injection at the source, normalized by the total number of pixels belonging to the target region. The tracer-based connectome is built by averaging over injection experiments performed in the right brain areas and targeting regions in ipsilateral and contralateral hemispheres. Through the Allen Connectivity Builder interface in TVB we parceled the brain in 104 anatomical regions of interest (ROIs, table in Table 1). Then, we defined the connection strength between source region *i* and target region *j* i.e., the link *ij*, as the ratio between the projection density at *j* and the injection density at *i*. The tracer connectome, with links normalized between 0 and 1 is shown in Figure 1c-d.

The empirical fMRI mice dataset was imported from the publicly available collection Grandjean (2020). For our analysis, we used the cohort of 20 control wild-type (resting) animals registered in the sub-dataset (Mandino, Yeow Ling Yun, Gigg, Olivo, & Joanes Grandjean, 2019). The data was previously preprocessed according to a common pipeline (https://github.com/grandjeanlab/MouseMRIPrep), and registered on the Allen volumetric atlas. In our analysis, we considered the activity in those voxels corresponding to the imported Allen connectome (in Table 1). The empiric dataset did not distinguish between specific parts (e.g.,ventral and dorsal) of certain brain regions (blue and red pairs in Table 1). A unique time series was associated with each of these pairs of regions.

The empirical EEG/fMRI dataset was acquired and preprocessed at Charit - Universittsmedizin Berlin (Schirner, McIntosh, Jirsa, Deco, & Ritter, 2018) and made available in online repository (https://osf.io/mndt8/) (Schirner, 2018). In summary, from a larger cohort (49 subjects, 18-80 years), 15 youngest subjects (18-31 years) were selected based on the quality of the EEG recording after correction of the MR artifacts. For each subject, diffusion-weighted MRI, T1-weighted MRI, and EEG-fMRI in resting-state paradigm were obtained. The T1-weighted image was used to parcellate the cortical gray matter into 68 regions according to the Desikan-Killiany atlas (Desikan et al., 2006). This definition of regions was then used to estimate the structural connectivity from the dw-MRI data (Schirner, Rothmeier, Jirsa, McIntosh, & Ritter, 2015), and to extract the regional average time series from the fMRI data. The EEG data (Easy-cap; 64 channels, MR compatible) was treated with a high pass filter at 1.0 Hz followed with MR imaging acquisition artifact removal using Analyser 2.0 (v2.0.2.5859, Brain Products, Gilching, Germany). The resulting sensor-level time series was downsampled to 200 Hz and low pass filtered to 30 Hz before correction for physiological artifacts (ballistocardiogram, muscle activity). Next, EEG source imaging was performed to obtain projected activity on the cortical surface and averaged within the 68 regions of the Desikan-Killiany parcellation. See Schirner et al. (2018) for a detailed description of both fMRI and EEG processing. After the preprocessing, we made a quality check of every subject data and excluded BOLD points and associated EEG time windows (BOLD TR = 1.94 s = 388 EEG samples) which presented residual artifacts (at either EEG or fMRI level) based on the following criteria:

- if any of the 6 time series of the motion degrees of freedom (resulting from the FSL MCFLIRT head movement correction step) presented a peak.
- if the EEG window contained fluctuations simultaneously affecting most of the frequencies (time-frequency analysis).
- if the EEG presented fluctuations above 7 standard deviations.

We finally selected 30 artifact-free trials of consecutive EEG/fMRI acquisition (minimum duration 2 minutes) across the cohort.

### 5.2 Neural Mass Model

The dynamics of a brain region of interest (i.e., a node in the network) is described by a Neural Mass Model (NMM) derived analytically as the limit of infinitely all-to-all coupled *θ*-neuron phase oscillators, whose dynamics is equivalent to that of quadratic integrate-and-fire (QIF) neurons (Figure 1a) (Montbrío et al., 2015). The *i*-th neuron is described by the equation

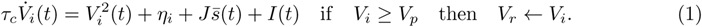

As the membrane potential reaches a peak value *V_p_*, it resets to *V_r_*. The limit *V_p_* = −*V_r_* → ∞ is considered. The parameter *η_i_* is the neuron excitability, and it enters as an heterogeneous current in the membrane potential. A time-dependent current *I*(*t*) homogeneously affects all the neurons in the neural mass. The synaptic activation *s*(*t*) of a single neuron respects the equation *Q · s(t) =Ʃ_k_δ(t − t_k_)*, being 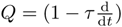 and *t_k_* the arrival time of the presynaptic action potential. The neuronal coupling among *N_QIF_* neurons inside a neural mass is given by the average synaptic activation 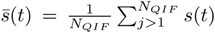 scaled by the synaptic weight *J*. Assuming a Lorentzian distribution of the membrane potentials across the coupled neuronal population it is possible to perform the *N_QIF_* → ∞ limit exactly according to the Ott-Antonsen ansatz (Ott & Antonsen, 2008). Also, the heterogeneous currents are distributed according to a Lorentzian distribution with width at mid-height Δ and peak location *η*. After the limit we can describe the activity in a neural mass *n* in terms of the average membrane potential *V_n_*(*t*) and the average firing rate *r_n_*(*t*) (which corresponds to 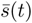 in the limit *τ* → 0). The dynamics associated with a local network node is then described by the following NMM equations

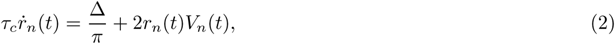

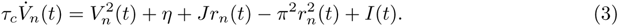

The synaptic weight *J* = 14.5, the average neuronal excitability *η* = −4.6, the spread of the heterogeneous noise distribution Δ = 0.7, and the characteristic time *τ_c_* = 1 are homogeneously tuned so that each decoupled node is in a bistable regime with a ‘down’ fixed point and an ‘up’ stable focus in the 2D phase space (Figure 1b, *I_ext_* = 0). Differently from standard NMMs, the chosen model keeps track of the internal amount of synchronization of the constituent neurons by preserving the analytic form of the Kuramoto parameter (Kuramoto, 2003)

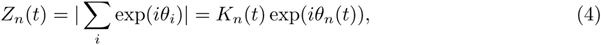

during the thermodynamic limit. The phase-oscillator representation (i.e., in terms of *θ_i_*) of the QIF equations can be achieved from (1) by a change of variables *V_i_* = tan(*θ_i_/*2). *K_n_* and *θ_n_* are the average amount of synchronization and the average phase of the neurons in the node *n*. The real and imaginary components of *Z_n_* are connected to the average membrane potential *V_n_* and firing rate *r_n_* of the neural population *n* via the relation

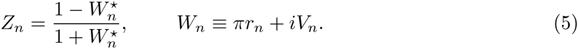

### 5.3 Connectome based large-scale Network Modeling

The local model is then coupled over an empirically extracted connectome (Figure 1d). The coupling term enters as an additive current in the average membrane potential equations (see also Figure 1e)

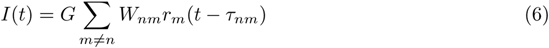

The global coupling parameter *G* sets the impact of the connectivity matrix *W_nm_* over the local dynamics. The time delay *τ_nm_* = *l_nm_/v* is given by the tract length between nodes *n* and *m* divided by the average conduction speed which we set to a realistic value *v* = 4 *m/s*. Since the firing rate is by definition greater than or equal to zero, all the interactions are excitatory.

The numeric integration of the nodal equations over the network is performed using The Virtual Brain (TVB) open-source neuroinformatics platform (Sanz Leon et al., 2013). The solution of the coupled system consists of a neuroelectric raw dataset describing the evolution of the variables (*r_n_*(*t*)*, V_n_*(*t*)) in each region *n* of the connectome. These variables are our measure of microscopic activity. The sampling rate of these neuroelectric variables is set to 1000Hz. The surrogate BOLD activity *B_n_*(*t*) in each region is derived by filtering the membrane potential with the Balloon-Windkessel model (Friston et al., 2000) (default values implemented in TVB). We use a repetition time of 2 seconds so that the BOLD rate is 0.5 Hz.

Note that the conduction speed *v* is a function of the physical tract lengths *l_nm_* of the empirical connectome and of the resolution *Dt* of the simulated signal, which physical interpretation is arbitrary. We impose it to be *Dt* = 1*ms*. Accordingly, 2000 time steps correspond to 2*s* = 1 BOLD point. All the dimensional arguments treated in the text are based upon this convention. In particular, we adopted a nondimensional formalism for the NMM equations (2), where only the characteristic time constant *τ_c_* has the dimension of time.

### 5.4 Time-dependent functional connectivity

To quantify the temporal evolution of the brain connectivity, we have employed two approaches: windowed dynamical functional connectivity (*dFC_w_*), and edge dynamical functional connectivity (*dFC_e_*). Let us denote *B_n_*(*t*) as the regional BOLD time-series for each node *n* = 1 … *N*. To compute the *dFC_w_* (Allen et al., 2014), we first obtain the series of functional connectivity matrices *FC*(*w*) at each sliding window *w* = 1 … *W*, defined as the correlation matrices for the segments 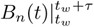 (in Figure 2b we fix the window size *τ* = 60*s* and window step 2*s*). Next, we correlate the vectorized upper triangular parts of the *FC*(*w*) matrices at different time windows to obtain the *dFC_w_* matrix

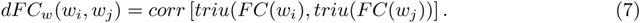

On the other hand, the computation of the *dFC_e_* starts with z-scoring the BOLD time series 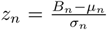. The edge time series is then computed as element wise multiplication along time for each pair of regions *E_nm_*(*t*) = *z_n_*(*t*)*z_m_*(*t*) for *n, m* = 1 … *N* (see Figure 1f). Next, for each pair of time points *t*_1_, *t*_2_ we compute the correlation of the edge vectors as

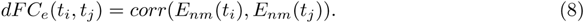

A main difference among the *dFC* variants lies in the scope of z-scoring of the time series *B_n_*. In the case of *dFC_e_* the z-score *z_n_* is computed from the whole time series, whereas in the *dFC_w_* the z-score is performed within the Pearson correlation in each time window [*t_w_, t_w_* + *τ*] separately.

### 5.5 Surrogate BOLD models

In order to compare our results in Figure 9a with the null hypotheses of a random evolution and of inter-regional stationarity of the *FC*, we build time-shuffled and phase-randomized surrogates of the *FC* dynamics, respectively. The time-shuffled surrogate is obtained by randomizing the order of the instantaneous functional connectivities *iFC*(*t*) i.e., the columns of the edge CA time series *E_ij_* (*t*) (see Figure 2a). According to Hindriks et al. (2016) the phase randomized surrogate is obtained by adding a uniformly distributed random phase to each frequency of the Fourier transformed signal, and subsequently retrieving the phase randomized signals by anti-Fourier transform. Importantly, the phase shift is different at every time point but can be applied uniformly to all the brain regions (cross-spectrum preserved) or separately in every region (cross-spectrum not preserved). Only in the first case the bursts of co-activation are not destroyed but shifted (Figure 9-1a) and the static *FC* is preserved. The coherent fluctuations around the stationary *FC*, however, are destroyed. For more details see also Prichard and Theiler (1994).

## Acknowledgments

The work was supported by the grants ANR-17-CE37-0001 - CONNECTOME and ANR-20-NEUC-0005-01 - Brainstim, by the European Unions Horizon 2020 research and innovation programme under grant agreement No. 945539 (SGA3) Human Brain Project and VirtualBrain-Cloud No. 826421.

## 7 Multimedia, Figure, and Table

**Figure 5-1:**
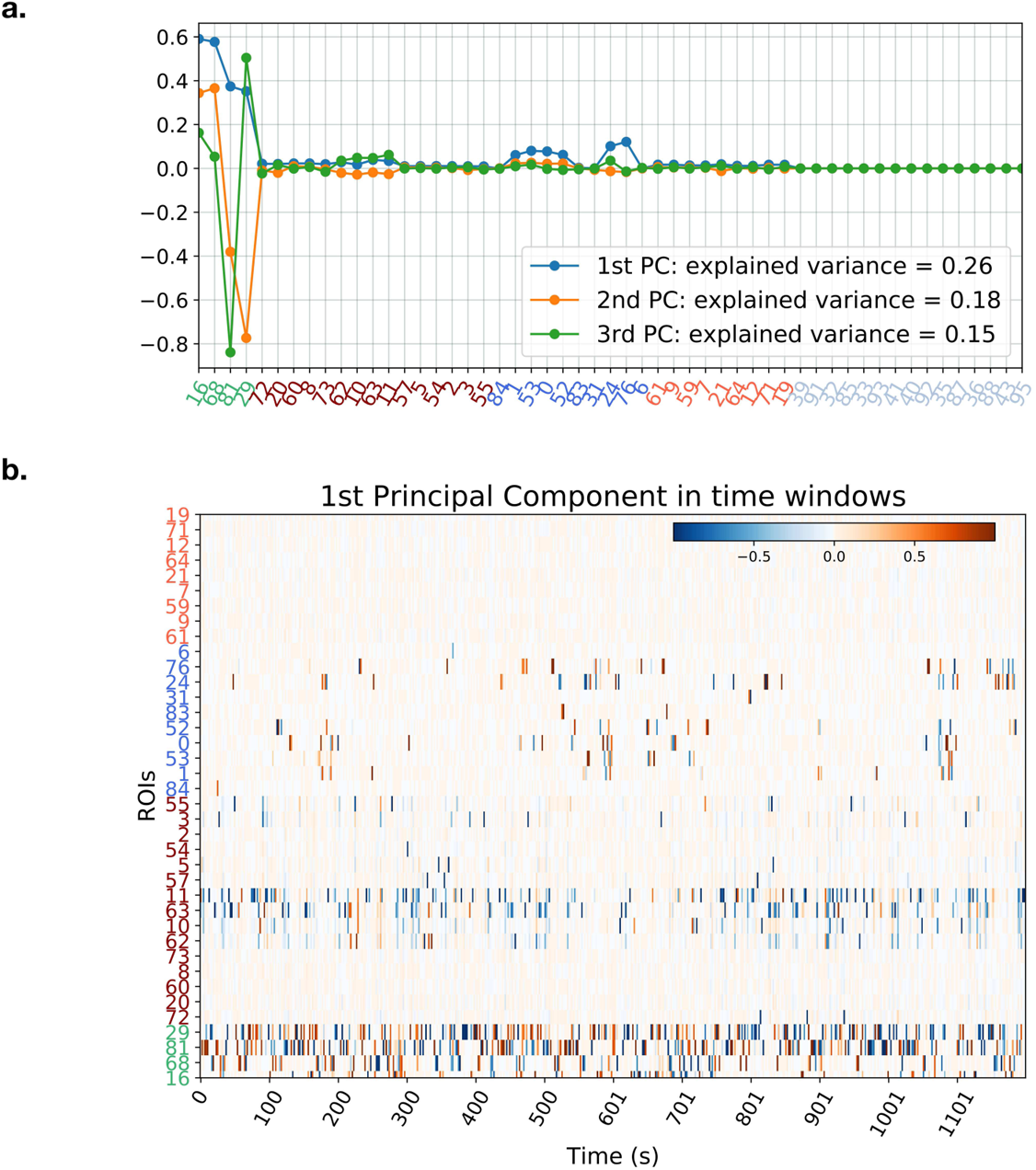
(a) The first three Principal Components extracted from 1200s of simulated firing rate activity in the bistable regime reveal a dominant contribution of the jumping regions (green ROIs corresponding to class J in Figure 4a). (b) The analysis of the first principal component of the simulated firing rate extracted from non-overlapping 2s time windows (corresponding to 2000 firing rate time points) reveals the intermittent contribution of the regions from classes U* and D* (dark red and dark blue ROIs, respectively), which become active carriers of the system variance during times associated to the neuronal cascades (e.g., around *t* = 600s corresponding to epoch III in Figure 5a).

**Figure 9-1:**
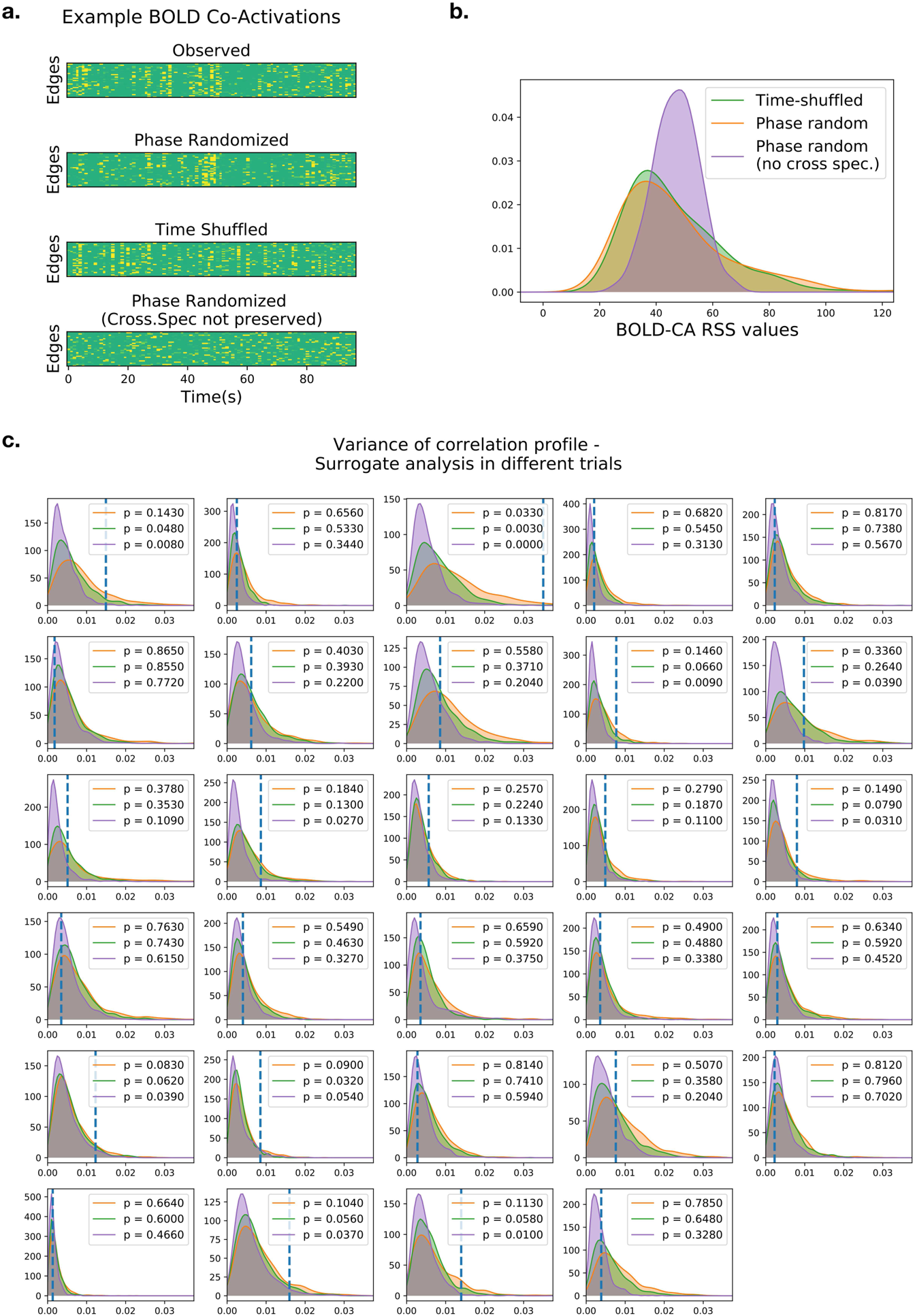
(a) Example of the edge CA time series in the original human dataset, compared to the respective phase-randomized and time-shuffled sur^5^r^8^ogates. Notice that the large co-activation events are preserved only in the first two surrogate models, but they are shifted in time. (b) Distribution of the BOLD-CA amplitudes (the root sum squared of the edge co-activations) in the time-shuffled (same distribution as in the original dataset) and in the phase randomized surrogates where the cross-spectrum is preserved (orange) or not (purple). (c) Single-trial analysis of variance distribution (similar to Figure 9b). The legend reports the p-values of the real observed variance versus the three surrogate distributions.

